# Why the middle ear piston prosthesis is not effective enough and how to change it?

**DOI:** 10.1101/2020.03.03.975052

**Authors:** Wiktor L. Gambin

## Abstract

Piston prostheses of the middle ear do not ensure sufficient audibility of high and low sounds. To find out the reason, the amplitudes of the vibrations for the stapes footplate and the piston end were compared. It was seen that for a given force that oscillates with a low frequency, the amplitude of the piston end was higher than the amplitude of the stapes footplate. This means that the stiffness of the tissue sealing up the piston in the footplate hole is lower than the stiffness of the stapes plate suspension. It was shown that as a result, for the case of the higher frequencies, the amplitude of the piston vibrations drops several times. Next, it was compared a sound propagation in the healthy ear with that in the ear after the stapedotomy. To do it, a previously prepared model of sound propagation in the ear was used. The model is simplified, but it gives all parameters of the sound wave in the cochlear fluid. According to it, a motion of the stapes footplate forms a plane wave, while the piston motion initially gives a wave similar to the spherical one. A part of the spherical wave with the front directed towards the apex forms the primary wave running in the cochlea. However, the rest of this wave has the front directed towards the stapes footplate. This wave part, after a reflection from the stapes footplate, creates a secondary wave that follows the primary wave. A splitting of the wavefront around the edge of the piston end is a source of disruptions in the sound perception. The shift of the secondary wave reduces the power of the primary wave; it disturbs the waving of the basilar membrane and may cause extra noise. To justify it, a graph of the level of the cochlear amplification for the ear with the piston prosthesis was shown. The result compared with a simulation for the healthy ear gave the values 5 dB lower. To remove these drawbacks, it was proposed to place the piston end, not inside the cochlea, but in a guide in the form of a tube ended with a funnel fixed in the hole made in the stapes footplate. The piston was suspended in a guide tube on an O-ring formed of silicone gel. It was shown that when the piston is in the guide, the level of cochlear amplification was the same as that in the healthy ear. Some design details of the new piston guide are given. It enables us to make the new piston prosthesis easily and put it into practice.

## 1. Introduction

In the late 50ies years, the concept of the piston prosthesis appeared [1]. Up to now, it is a commonly used a passive implant of the middle ear [2]. Despite the undoubted improvement of hearing of sounds in the range of the middle frequencies (the speech sounds), the high and the low sounds are hard to hear [3]. It was noted that some patients complain of echo-like noise heard in the first month after the stapedotomy.

To explain it, a change of the amplitude of the vibrations with their frequency for the stapes footplate and the piston end was compared. In both cases, the same force was taken to get low-frequency vibrations. In that case, the amplitude of the piston end was twice higher than the amplitude of the stapes footplate. It was assumed that the stiffness of the tissue sealing up the piston in the footplate hole is twice lower than the stiffness of the stapes plate suspension. With this assumption, it was shown that in the case of the higher frequencies the piston amplitude drops several times.

Next, the shape of the wavefront which propagates in the cochlea of the healthy ear with that in the ear after the stapedotomy was compared. To do it, a simplified model of sound propagation in the human ear was used [4]. The model has enabled us to get all parameters of the sound wave in the cochlear fluid. One of its assumptions was to omit the rocking-like motions of the stapes footplate that occur at sound frequencies greater than 2.5 kHz. Despite this assumption, the model gives a good estimation of the sound wave pressure inside the cochlea up to 10 kHz. According to it, a motion of the stapes footplate of the healthy ear forms a plane wave in the cochlear fluid, but the end of thin piston initially creates a wave close to a spherical one. At a distance from the piston end, this wave becomes a flat one due to reflections from the cochlea walls. Close to the piston end, only a part of the spherical wavefront is directed towards the cochlea apex. The part remain of the wavefront is directed towards the stapes footplate. The first part forms a **primary wave**. The second part of the wavefront, after a reflection from the fixed stapes footplate, creates a **secondary wave**. The secondary wave follows the primary one with some delay. The reduced power of the primary wave and the shift of the secondary wave disturb a motion of the basilar membrane. This leads to a decrease in sound perception and makes echo-like noise. One can show the effect of the wave split on a graph of the level of the cochlear amplification function (see [5]). This graph for the ear with the piston prosthesis compared with that for the healthy ear gives values 5 dB lower.

To remove these obstacles, it was proposed to place the piston end, not inside the cochlea, but in a guide tube in the form of pipe ended with a funnel fixed in the hole made in the stapes footplate. This may allow producing in the cochlea only a flat wave focused in the outlet of the guide. The piston can be suspended in a guide tube on an O-ring formed of the silicone gel. Its stiffness should be the same as the stiffness of the annular ligament. This will ensure that in the case of the higher frequencies, a drop of the amplitude of the piston vibrations will not occur.

This work aims to analyze the flow of the sound wave in the cochlea with piston prosthesis. This aim includes a rating of the effectiveness of the piston prosthesis, which is free from the disadvantages of the currently used ones. Shown in the work, design details of the new piston prosthesis enable us to make it and put it into practice.

## 2. The reduced natural frequency of the piston prosthesis

Now, it will be shown the impact of the piston static amplitude growth on a drop its dynamic amplitude. Assume that the sound reaches the ear with a fixed sound intensity level ***β*** = 90 [*dB*]. Then, according to the work [4], on the stapes footplate acts a harmonic force with the amplitude ***N***_***sf***_ = 1.4052 ⋅ 10^−6^ [*N*]. The same force acts on the piston when the same sound reaches the ear with the piston prosthesis. Let us denote the measured amplitudes of the stapes footplate and the piston end by ***d***_***sf***_ and ***d***_***p***_. The test results for the frequencies 0.4 *kHz* ≤ ***f***_(***i***)_ ≤ 1.0 *kHz*, are given in the work [6] and shown in Table 1.

**Table 1.**

| *Test*<br>*i* | *Frequency*<br>*$f_{(i)}: [Hz]$* | *Measured amplitude*<br>*$d_{sf} [mm]$* | *Measured*<br>*amplitude*<br>*$d_p [mm]$* | *Amplitude ratio*<br>*$\frac{d_p}{d_{sf}}$* |
| --- | --- | --- | --- | --- |
| 1 | 400 | $1.64 \cdot 10^{-05}$ | $2.93 \cdot 10^{-05}$ | 1.8 |
| 2 | 500 | $1.53 \cdot 10^{-05}$ | $2.62 \cdot 10^{-05}$ | 1.7 |
| 3 | 630 | $1.90 \cdot 10^{-05}$ | $1.62 \cdot 10^{-05}$ | 0.9 |
| 4 | 800 | $1.97 \cdot 10^{-05}$ | $1.89 \cdot 10^{-05}$ | 1.0 |
| 5 | 1000 | $1.02 \cdot 10^{-05}$ | $1.05 \cdot 10^{-05}$ | 1.0 |
| 6 | 1250 | $8.92 \cdot 10^{-06}$ | $7.31 \cdot 10^{-06}$ | 0.8 |
| 7 | 1600 | $7.12 \cdot 10^{-06}$ | $8.58 \cdot 10^{-06}$ | 1.2 |
| 8 | 2000 | $4.93 \cdot 10^{-06}$ | $6.25 \cdot 10^{-06}$ | 1.3 |
| 9 | 2500 | $2.84 \cdot 10^{-06}$ | $3.48 \cdot 10^{-06}$ | 1.2 |
| 10 | 3150 | $6.40 \cdot 10^{-07}$ | $9.10 \cdot 10^{-07}$ | 1.4 |
| 11 | 4000 | $7.73 \cdot 10^{-07}$ | $3.35 \cdot 10^{-07}$ | 0.4 |
| 12 | 5000 | $8.36 \cdot 10^{-07}$ | $2.62 \cdot 10^{-07}$ | 0.3 |
| 13 | 6300 | $6.14 \cdot 10^{-07}$ | $1.59 \cdot 10^{-07}$ | 0.3 |
| 14 | 8000 | $1.69 \cdot 10^{-07}$ | $1.43 \cdot 10^{-07}$ | 0.8 |
| 15 | 10000 | $7.57 \cdot 10^{-7}$ | $1.85 \cdot 10^{-07}$ | 0.2 |

One can see that for the low frequencies when ***f***_(***i***)_ ≤ 0.5 *kHz* the amplitude of the piston end is higher than the amplitude of the stapes footplate. Whereas, for high frequencies when ***f***_(***i***)_ ≥ 4.0 *kHz*, it is the opposite.

In the work [4], for the given stiffness of the annular ligament 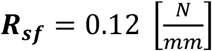, the static amplitude of the stapes footplate 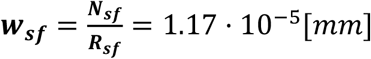 is found. This value is close to the measured ***d***_***sf***_ = 10.2 *nm* for the sound frequency ***f***^∗^_(***i***)_ = **1000** ***Hz***. It is the ***natural frequency*** of the stapes footplate, for which in the absence of damping a resonance should take place.

The static amplitude of the piston ***w***_***p***_ is unknown. To find its impact on the dynamic amplitude, one can assume, for example, that it is twice that for stapes footplate. Let us take ***w***_***p***_ = **2*****w***_***sf***_ = 2.34 ⋅ 10^−5^[*mm*] which corresponds with the stiffness of the piston suspension 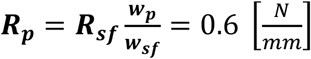. From Table 1, it yields that the value ***w***_***p***_ = 2.34 ⋅ 10^−5^[*mm*] is close to the measured ***d***_***p***_ = 26.2 *nm* for the sound frequency ***f***^∗∗^_(***i***)_ = **500** ***Hz***, which can be taken as the natural frequency of the piston. Introducing for the piston the ***resonant angular frequency*** 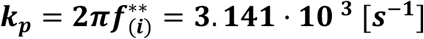 and the damping factor **2*****n***_***p***_ = ***k***_***p***_ (see [4]), one can find its dynamic amplitude ***A***_***p***_ (see Eq. (21) in [4])

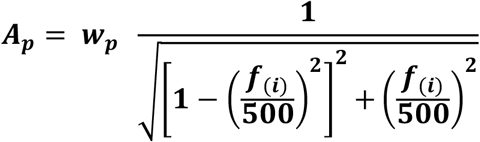

The comparison of the calculated values of the dynamic amplitude ***A***_***p***_ with the measured ones is given in Table 2.

**Table 2.**
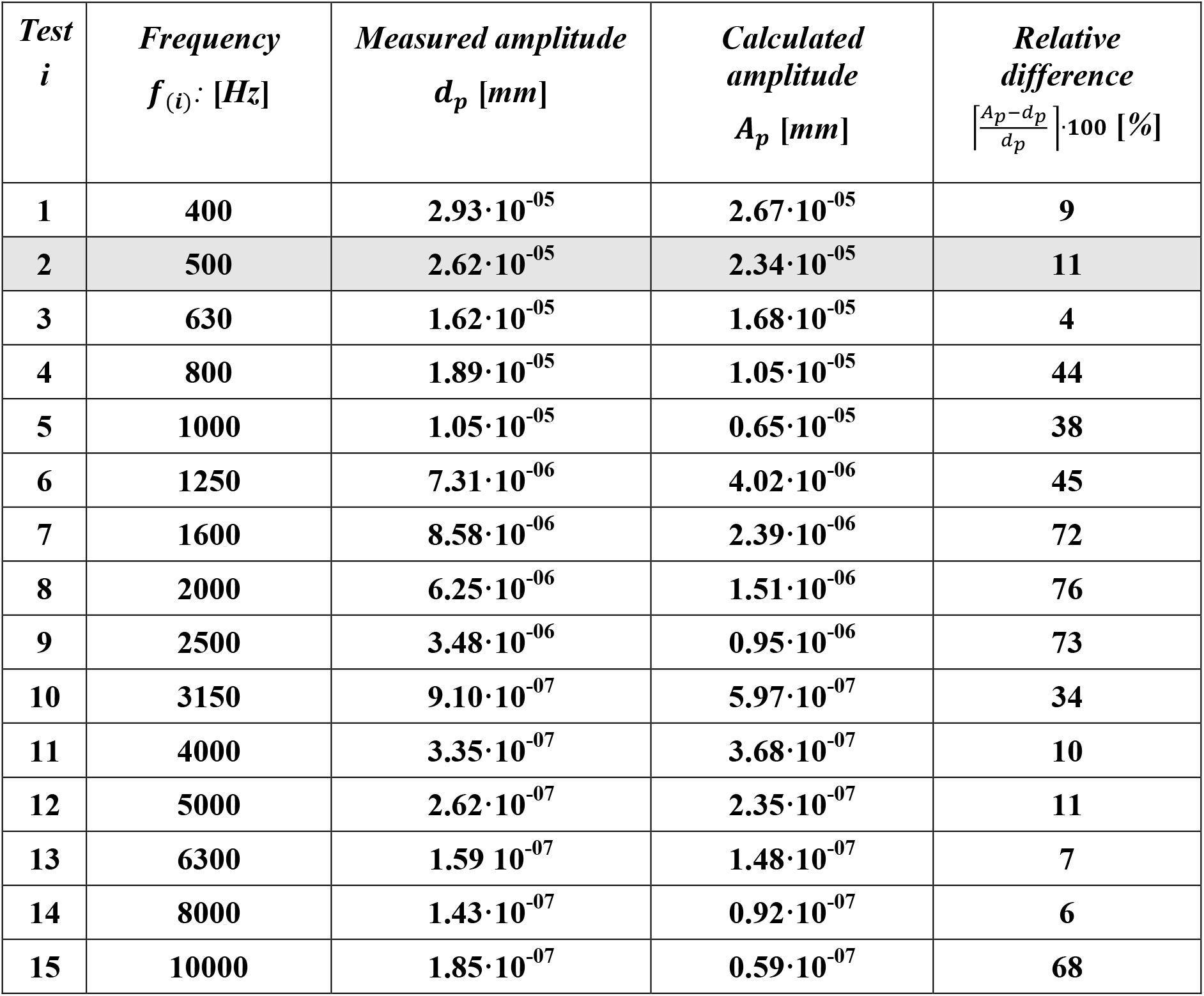

### Consequence

Due to the twice smaller stiffness of the piston suspension, the piston has the twice smaller natural frequency. It makes a large drop of the piston end amplitude for the frequencies 4000 Hz and the higher. To avoid this, one should change the piston suspension.

## 3. Is the sound wave that runs above the basilar membrane flat?

### Basic assumptions

At first, recall the basic assumptions of the model presented in the work [4]. Let from a sound source runs in the air a sound wave at the given level of sound intensity ***β*** and with a frequency ***f***_(***i***)_. Then, one can assume.

1. The outer ear is treated as a straight-line waveguide with a changing cross-section, filled with air and ended with an ***elastic*** tympanic membrane. In the waveguide runs an ***elastic plane*** wave.
2. The reduction of the sound intensity caused by a reflection of the sound wave from the tympanic membrane is due to only the difference between the wave resistance of the air ***Z***_***0***_ in the ear canal and the wave resistance of the fluid ***Z***_***1***_ in the cochlea.
3. The middle ear contains a lever system formed by a set of flexibly joined, stiff ossicles with a stapes footplate suspended on the ***viscoelastic*** annular ligament.
4. The inner ear is treated as a waveguide of changing cross-section filled with fluid, which runs an ***elastic plane wave*** forced by vibrations of the stapes footplate. Its length **2*L***_***c***_ is a double distance from the oval window to the cochlea apex measured along the spiral axis of the cochlea. The axis of this waveguide is straight, but in the middle of the waveguide, it changes direction to the opposite one. In this way, the waveguide is divided into two parallel-connected parts, but which direct the sound wave in the opposite directions.
5. Between these two parts lays, a chain of ***uncoupled viscoelastic fibers*** of length ***L***_***bm***_, which oscillate under the pressure of the sound wave running in the cochlear fluid. The chain starts at a distance ***x***_**0**_ from the oval window. Mass, stiffness, and viscosity of each of the fibers are known.
6. A waving of the chain of the fibers does not influence on the amplitude of the running sound wave.
7. The outlet of the waveguide with the set of fibers is closed by an ***elastic*** membrane of the round window, which fully absorbs the energy of the running sound wave.

The above assumptions enabled to find:

- a pressure and amplitude of the sound wave in the outer ear,
- the force acting on the stapes footplate, and the amplitude of the forced vibrations,
- a pressure and amplitude of the sound wave in the inner ear,
- the amplitude of the fibers vibrations due to the running sound wave,
- the amplitude of the vibrations of the round window membrane.

The results based on this simplified model have appeared to be quite close to the test results made with help of the temporal bone specimens [5–7].

### A flattening of the sound wavefront

It was shown in [4], that the sound intensity on the front of the sound wave in the cochlear fluid is inversely proportional to the cochlear cross-section. One can see in Fig. 1, the part of the cochlear waveguide with the scala vestibule and the scala tympani. Let us look on the shape of the front of the sound wave generated by the vibration of the stapes footplate due to a harmonic force ***N***_***sf***_ (***t***). According to the Huygens principle and the taken assumptions, a flat surface of the vibrating stapes footplate makes a flat wave, but its edge makes a deflection of the wavefront. The cross-section of the bent part becomes the arc of a circle. Thus, a surface area ***F***_***wf***_ (***x***) of the wavefront, at a distance ***x*** from the oval window, is slightly larger than the cross-sectional area ***F***_***c***_ (***x***) of the cochlea channels. Let ***I***_***c***_ (***x***) is the intensity of the sound wave at the point ***x***. Because of the sound power ***I***_***c***_(***x***) ⋅ ***F***_***c***_ (***x***) is fixed, the increase of the area of the wavefront from ***F***_***c***_ (***x***) to ***F***_***wf***_ (***x***), suggests that ***I***_***c***_ (***x***) is lower. On the other hand, due to wavefront reflections from the walls of the cochlea, this deflection is fast reduced. Let us try to estimate the impact of both factors.

**Fig. 1.**
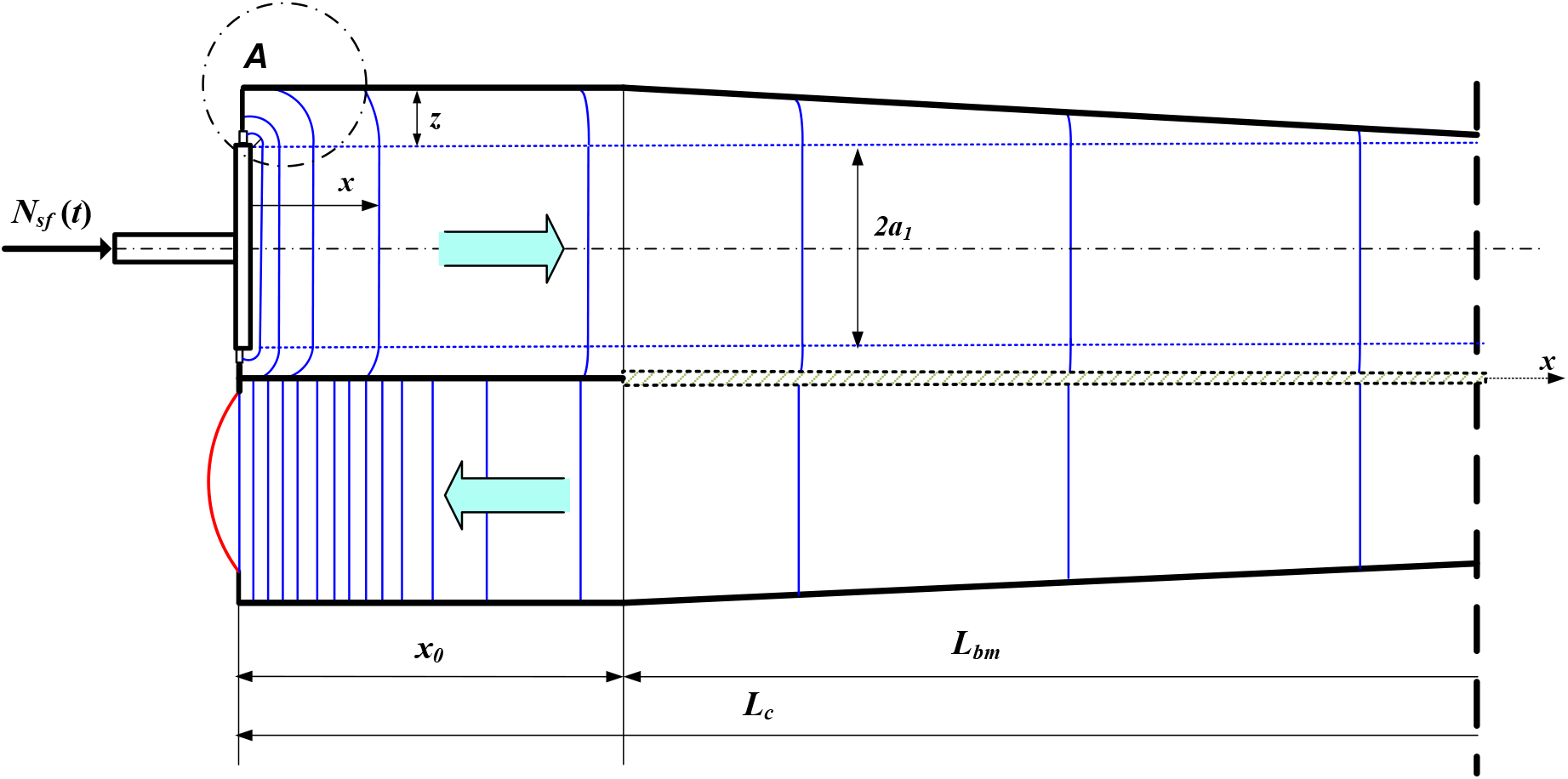
Wavefront generated by vibrating stapes footplate.

Let us try to estimate the rate of the reduction of the wavefront deflection. To simplify, let us take that the cross-sections of the cochlea between the oval window and the beginning of the basilar membrane are the same. Denote by ***z*** a distance between the edge of the stapes footplate and the nearest cochlea wall. Consider the element ***A*** in Fig. 1 in which forms the curved front wave of the sound wave.

Fig. 2 shows phases of the wave formation reflected from the cochlear walls. When the wavefront crosses the point ***x*** = ***z***, the wave reflection appears. When it reaches the point 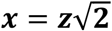, the power of the output wave and the wave reflected from the wall are balanced. At that time, the resultant vector of the momentum of both waves is parallel to the ***x***-axis. Finally, when the output wavefront gets the point ***x*** = **2*****z***, the reflected wave reaches the line separating the flat and the curved part of the output wavefront. One can assume that starting from the cross-section ***x*** = **2*****z***, the whole front of the output wave is flat.

**Fig. 2.**
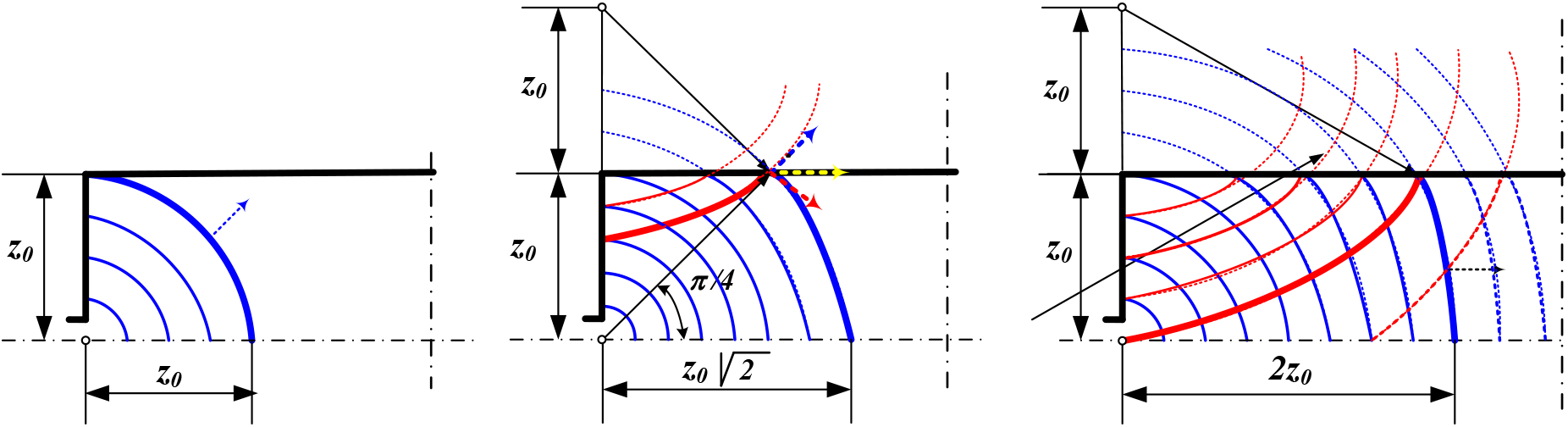
Flat wavefront formation as a result of wave reflection from the cochlear wall.

To find the distance ***z*** between the edge of the stapes footplate and the nearest cochlea wall, the shape, and dimensions of the cochlear cross-section for ***x*** = ***x***_**0**_ = 2.6 *mm* are given in Fig. 3a (according to [8]). For the cochlea with the piston prosthesis, one can take the distance ***z***^***p***^ between the edge of the piston end and the nearest cochlea wall (Fig 3b).

**Fig. 3.**
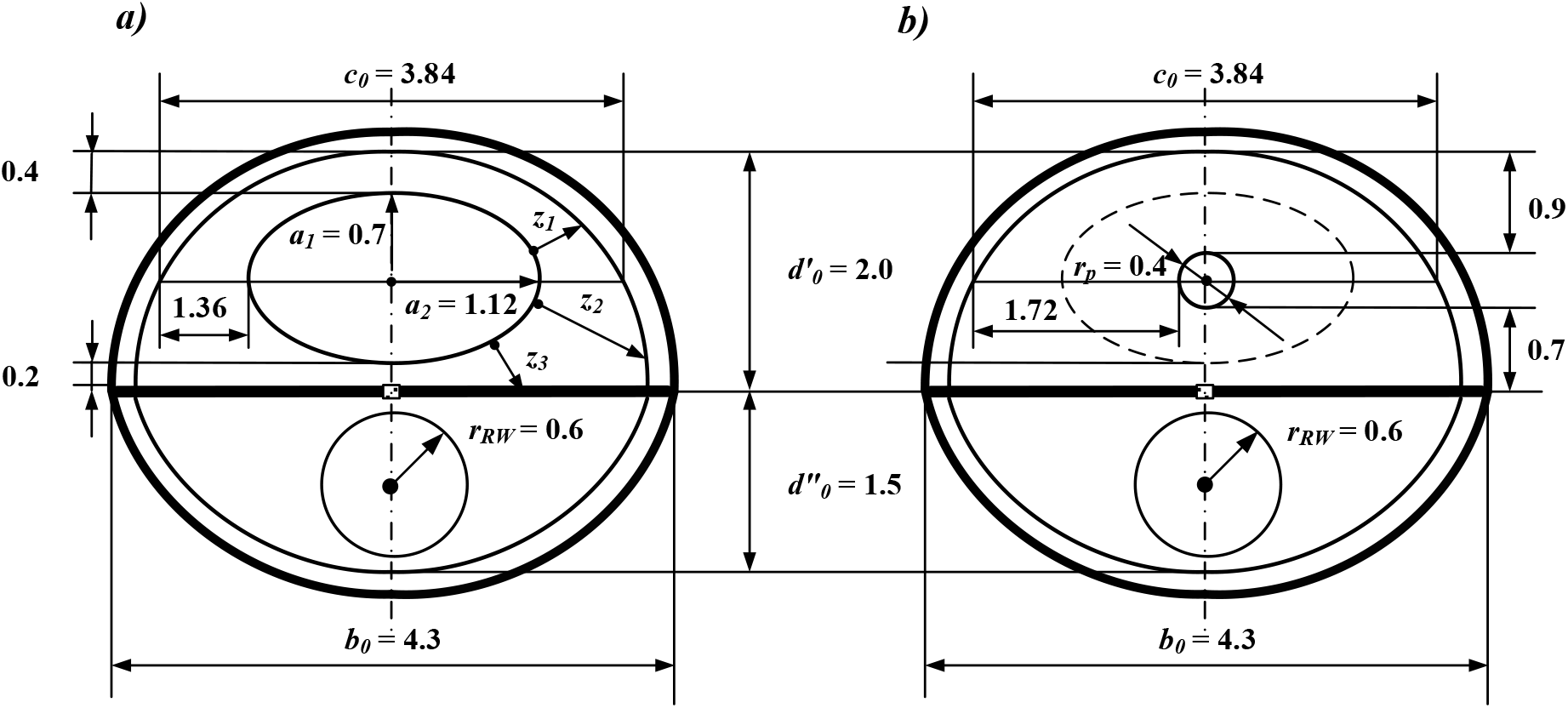
Shapes and dimensions in mm of the cochlear cross-section at point ***x*** = ***x***_**0**_, for the healthy ear (based on [8]) (a) and for the ear with piston (b).

The cross-section of the vestibule channel has the shape of half ellipse with the semi-axis ***d***′_***o***_ = 2.0 *mm* and the axis ***b***_***o***_ = 4.3 *mm*. The stapes footplate is taken as an ellipse with semi-axes ***a***_**1**_ = 0.7 *mm* and ***a***_**2**_ = 1.12*mm*. A diameter of the piston is equal to **2*****r***_***p***_ = 0.4 *mm*.

Fig. 3 shows that the distance ***z*** varies within the cross-sections. We can use instead of ***z***, its average values ***z***_***aν***_. To do it, let us introduce a width of the vestibule channel at the height of the center of the stapes footplate equal to ***c***_**0**_ = 3.84 *mm*. Now, one can take, for the oscillating stapes footplate

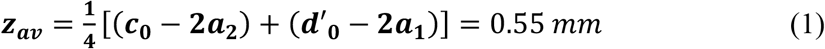

and for the vibrating piston

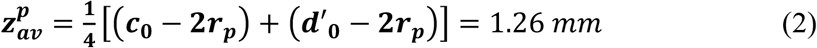

Note that the distance **2*****z***_***aν***_ = 1.1 *mm* < ***x***_**0**_ = 2.6 *mm*. Same, the distance 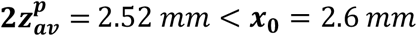. So, at the base of the basilar membrane, the front of the sound wave, both in the healthy ear and the ear with piston prosthesis can be taken as a flat one.

## 4. A secondary sound wave in the cochlea with the piston prosthesis

### The taken assumptions

Fig. 4 shows a cochlear waveguide of the same shape and dimensions as presented in Fig.1. In the hole made in the stapes footplate, a piston of diameter **2*****r***_***p***_ = 0.4 *mm* is placed to a depth of ***l***_**0**_ = 1.0 *mm*. The cross-sectional dimensions of the cochlea with a piston for ***x*** = ***x***_**0**_ are given in Fig. 3b. A length ***z***^***p***^ is a distance between the edge of the stapes footplate and the nearest cochlea wall, and 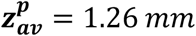 is its average value at the cross-section of the cochlea.

**Fig. 4.**
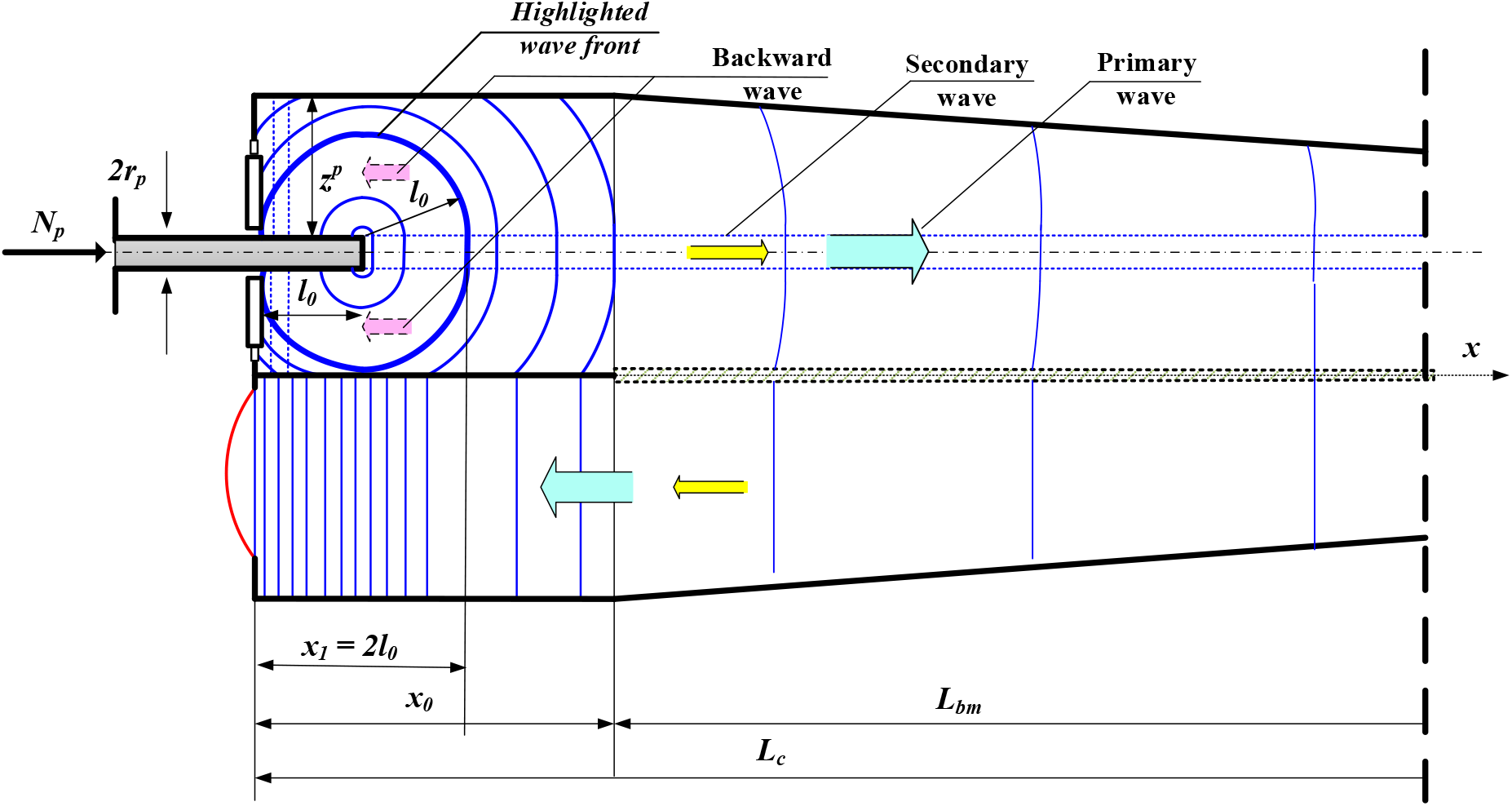
Wavefronts generated by vibrating piston prosthesis.

### The effect of the sound wave duplication

Now, around the end of the piston, a sound wave similar to a spherical wave is seen. One can expect, the sound wave intensity ***I***_***cp***_(***x***) in the cochlea with an implanted piston, will change concerning the value ***I***_***c***_(***x***). To find this change, assume that a sound wave of intensity ***β*** and frequency ***f***_(***i***)_ reaches the ear with the implant. Then the force ***N***_***p***_ acting on the piston is the same as the force ***N***_***sf***_ acting on the stapes footplate in a healthy ear. According to our model, the power ***P***_***cp***_ of the sound wave in the cochlea with the piston is equal to the piston power ***P***_***p***_. The total power ***P***_***cp***_ is transmitted by those wavefronts that do not have contact with the cochlea walls or the stapes footplate. For further analysis, we choose from theses wavefronts that one which is furthest from the end of the piston (see Fig.4). Its distance from the piston end is the smaller of two values: ***z***^***p***^ and ***l***_**0**_.

In our example, this value is the distance ***l***_**0**_ < ***z***^***p***^. So, the position of the highlighted wavefront on the ***x***-axis is fixed by the coordinate ***x***_**1**_ = **2*****l***_**0**_ = 2.0 *mm*. Now, the power ***P***_***cp***_ can be expressed by the sound wave intensity ***I***_***cp***_(***x***_**1**_) at the highlighted wavefront

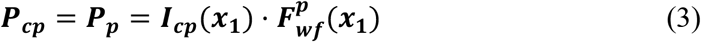

where 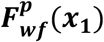 is its surface area.

Looking at Fig. 4, and next in Fig.5, one can see that the surface of the highlighted wavefront can be decomposed onto two parts. The first of them has normal vectors ***n*** inclined towards the apex of the cochlea, and the second one ‒ towards the stapes footplate. The first surface consists of two elements. The first element is a flat circle with a radius ***r***_***p***_. The second element is a quarter of the spindle torus surface. The major radius of the torus is equal to ***R*** = ***r***_***p***_, and its minor radius is ***r*** = ***l***_**0**_.

**Fig. 5.**
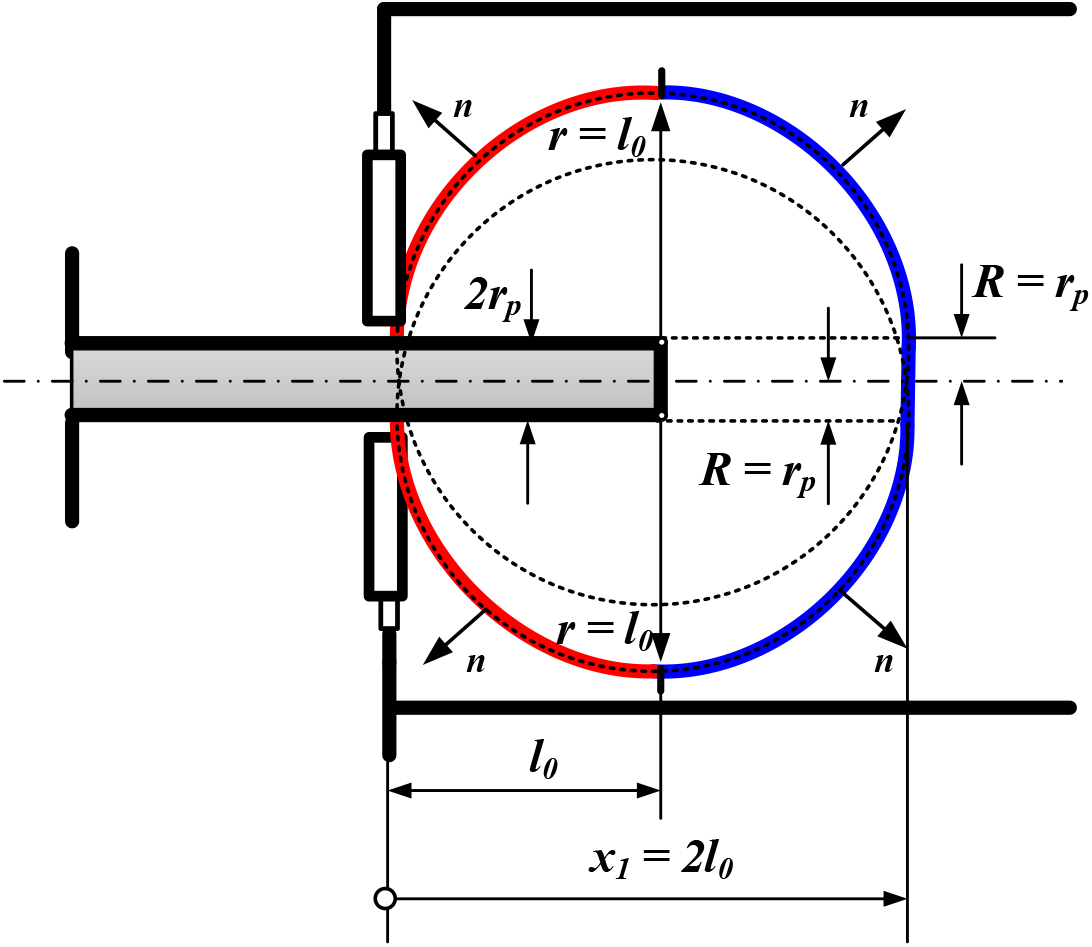
Decomposition of highlighted wavefront formed by piston end.

Area of the first surface of the highlighted wave front-facing to the apex of the cochlea is

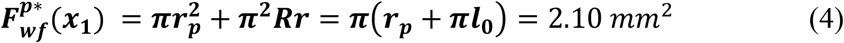

The second part of the highlighted wavefront, facing to the stapes footplate, is a quarter of the spindle torus surface with the area

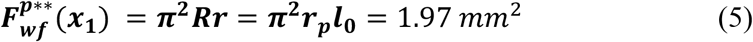

The total surface area of the highlighted wavefront is

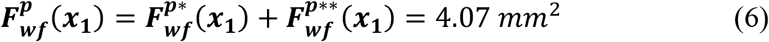

The first part of the wavefront forms a front of the wave called a **primary wave**. The second part, directed towards the stapes footplate gives a **backward wave**. The backward wave after the reflection from the fixed stapes footplate forms a **secondary wave.** The secondary wave follows the primary wave at a short distance which is equal to **2*****l***_**0**_. At a distance of ***l***_**0**_ from the end of the piston, both wavefronts can be taken as flat. Note that the power ***P***_***cp***_ can be decomposed as follows

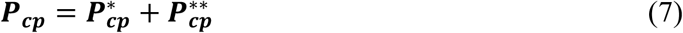

where

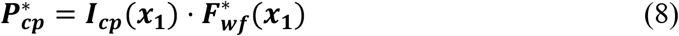

is a power of the primary wave, and

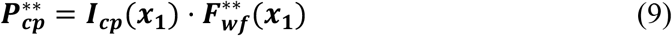

is the power of the backward wave equal to the power of the secondary wave. Let us look at the share of the secondary wave on the process of the sound propagation in the cochlea. Because ***P***_***cp***_ = ***P***_***p***_, for the dimensions shown in Fig. 3, one can get

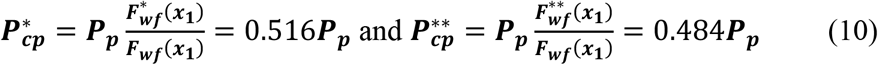

Note that only half of the sound power reaching the ear is used by the primary wave. Also, the relative shift and delay of the wavefronts can be relevant. Here, the shift is **2*****l***_**0**_ = 2.0 *mm*, and the delay is **2*****l***_**0**_/***c***_**1**_ = 1,3699 ⋅ 10^−6^ *s*, where ***c***_**1**_ = 1.46 ⋅ 10^6^ *mm*/*s* is the speed of the sound wave in the cochlea. The delay is very small, but it can interfere with the hearing process. Surely the part of the sound power which forms the secondary wave disturbs a run of the primary wave.

## 5. Impact of the sound wave splitting on the basilar membrane amplitude

### The basilar membrane amplitude in a healthy ear

According to the model given in [4], the ***x*****-**axis runs along the vestibular duct to the apex, then turns back and it runs along the tympanic duct to the round window (Fig. 6). At a distance ***x***_**0**_ = 2.6 *mm* from the oval window, a basilar membrane with a length ***L***_***bm***_ = 31.9 *mm* starts. It runs along the ***y*****-**axis up to the apex. The piston end inserted into the cochlea to a depth of ***l***_**0**_ = 1.0 *mm*.

**Fig. 6.**
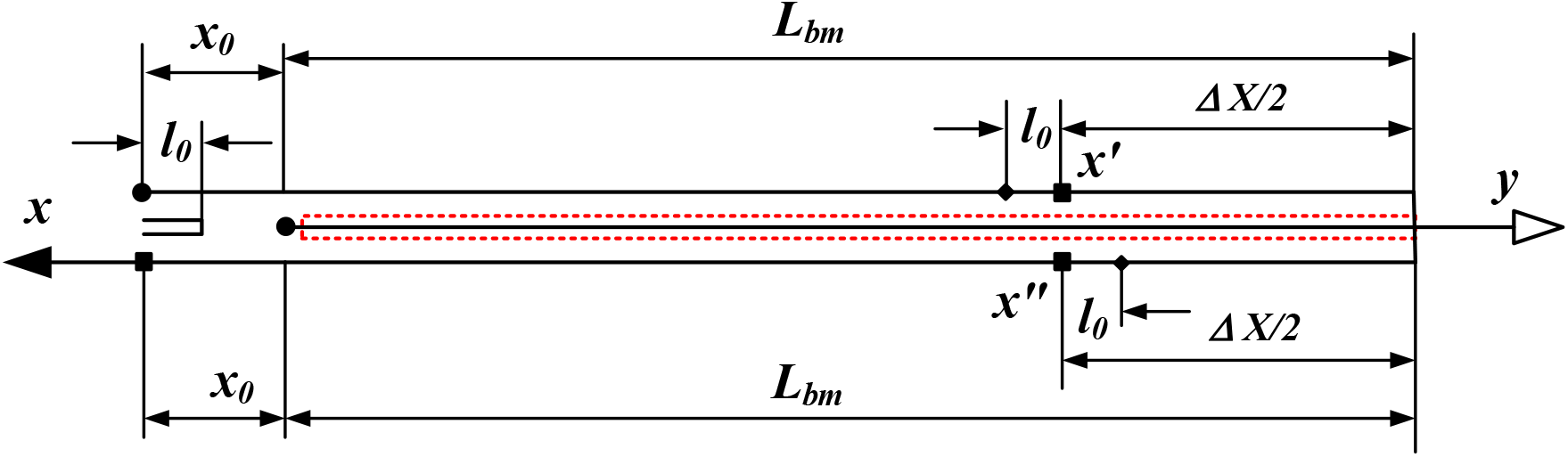
Model of the basilar membrane.

Let us consider a point ***y*** of the basilar membrane. This point has a coordinate ***x***′ in the vestibular duct and a coordinate ***x***" in the tympanic duct.

The relationships between the coordinates ***x***′, ***x***" and ***y*** have the form

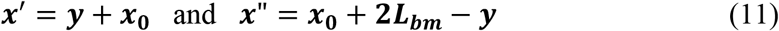

In a healthy ear, each point of the basal membrane is pressed by a sound wave with a frequency ***f***_(***i***)_ and the amplitude ***A***_***c***_(***x***). This wave flows both in the vestibular duct and in the tympanic duct. The sound wave pressure forms on the basilar membrane a wave with the amplitude ***A***_***bm***_(***y***) that runs along the membrane back and forth. It was assumed in the work [1], that the amplitude ***A***_***bm***_(***y***) is due a difference **Δ*****p***_***c***_(***y***) between the sound pressures

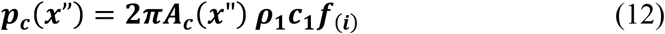

and

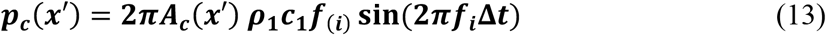

where ***c***_**1**_ = 1.46 10^6^ *mm*/*s* is the sound velocity in the cochlea fluid and ***ρ***_**1**_ = 1.0 ∙ 10^−3^ *g*/*mm*^3^ is its density. Here, ∆***t*** = **2**(***L***_***BM***_ − ***y***)/***c***_**1**_ is the delay of the wavefront at the point ***x***" concerning that at the point ***x***". One can show that (see [4])

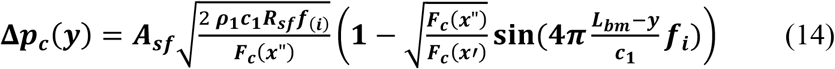

where ***A***_***sf***_ is the amplitude of the stapes footplate, ***R***_***sf***_ ‒ the stiffness of the annular ligament, ***F***_***c***_(***x***”) and ***F***_***c***_(***x***”) ‒ the areas of the cochlear cross-sections.

It was shown in the work [4], the amplitude ***A***_***bm***_(***y***) is expressed by **Δ*****p***_***c***_(***y***) by the rule

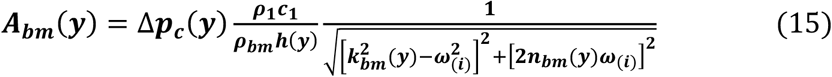

where ***ρ***_***bm***_ = 1.20 ⋅ 10^−3^ *g*/*mm*^3^ is the basilar membrane density, and ***h***(***y***), ***k***_***bm***_(***y***) and ***n***_***bm***_(***y***) are the thickness, the resonant frequency and damping coefficient of the basilar membrane at the point ***y***. It yields from Eqs (14–15) that ***A***_***bm***_(***y***) is uniquely given by the stapes footplate amplitude ***A***_***sf***_.

### The basilar membrane amplitude in the ear with the piston prosthesis

Recall that the assumed stiffness of the piston suspension is 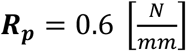, and the amplitudes of the piston end ***A***_***p***_ for 0.4 *kHz* ≤ ***f***_(***i***)_ ≤ 1.0 *kHz* are given in Table 2. According to Eq. (10), the power of the primary wave 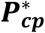 is only 0.516 of the piston power ***P***_***p***_. At a fixed point ***x***, the pressure of the primary wave 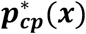 is proportional to the square of the power 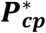. Thus, changing ***A*** (***x***") on 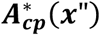 and ***A*** (***x***′) on 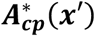 in Eqs (12–13), we have 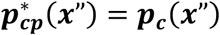 and 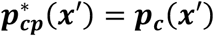. From Eq. (14), one can get the pressure difference 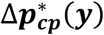 due to the **primary wave**

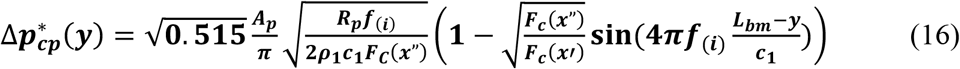

Now, let us find the pressure difference 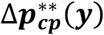 due to the secondary wave. When the primary wavefront is in the tympanic duct at the point ***x***", the secondary wavefront will appear at this point after the time ∆"***t*** = **2*****l***_**0**_/***c***_**1**_ = 1,3699 ⋅ 10^−6^ *s* (see Fig. 6). Thus,

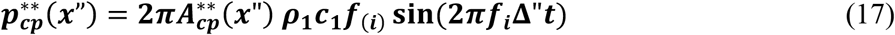

At the same time, the secondary wavefront in the vestibular duct, at the point ***x***′, is delayed concerning the primary wavefront, at the point ***x***", by ∆′***t*** = **2**(***l***_**0**_ + ***L***_***BM***_ − ***y***)/***c***_**1**_.

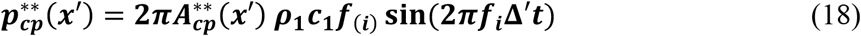

Thus, we get the pressure difference 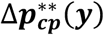 due to the **secondary wave**

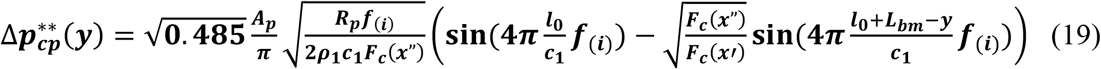

Since the secondary wave follows the primary one, both give the pressure difference ∆***p***_***cp***_(***y***) equal to

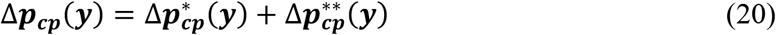

Concluding, the amplitude of the basilar membrane in the ear with a piston prosthesis is given by the rule

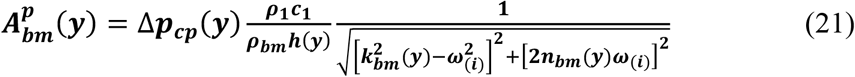

Looking at Eq. (16) and Eq. (19), one can see that for a given the piston amplitude ***A***_***p***_.the amplitude ***A***_***bm***_(***y***) is known.

### Level of cochlear amplification

Let us consider a fixed point ***y***^∗^ of the basilar membrane. Its coordinate in the tympanic duct given is ***x***^∗^ = **2*****L***_***bm***_ + ***x***_**0**_ − ***y***^∗^. At this point, the amplitude 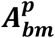 can be shown as a function of the sound wave frequency ***f***_(***i***)_ and the parameter ***y***^∗^. The function

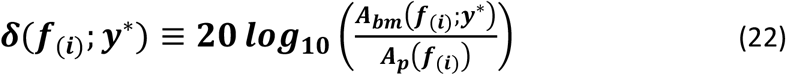

shows a ***level of cochlear amplification*** at the point ***y***^∗^.

The results of measurements of the function ***δ***(***f***_(***i***)_; ***y***^∗^) are given in the work [5]. The tests were done at the point located at ***y***^∗^ + ***x***_**0**_ = **12** [*mm*] from the oval window for different sound wave frequencies. The results were set in ***dB*** and normalized with the velocity of the stapes footplate. On the base of the rules (16-22) and the assumed data, one can plot the function ***δ***(***f***_(***i***)_; ***y***^∗^). Fig 7 shows a comparison of the test results with estimations made for a healthy ear and for the ear with the piston prosthesis. One can see a 5 dB drop in ear-prosthesis results compared to a healthy ear.

**Fig. 7.**
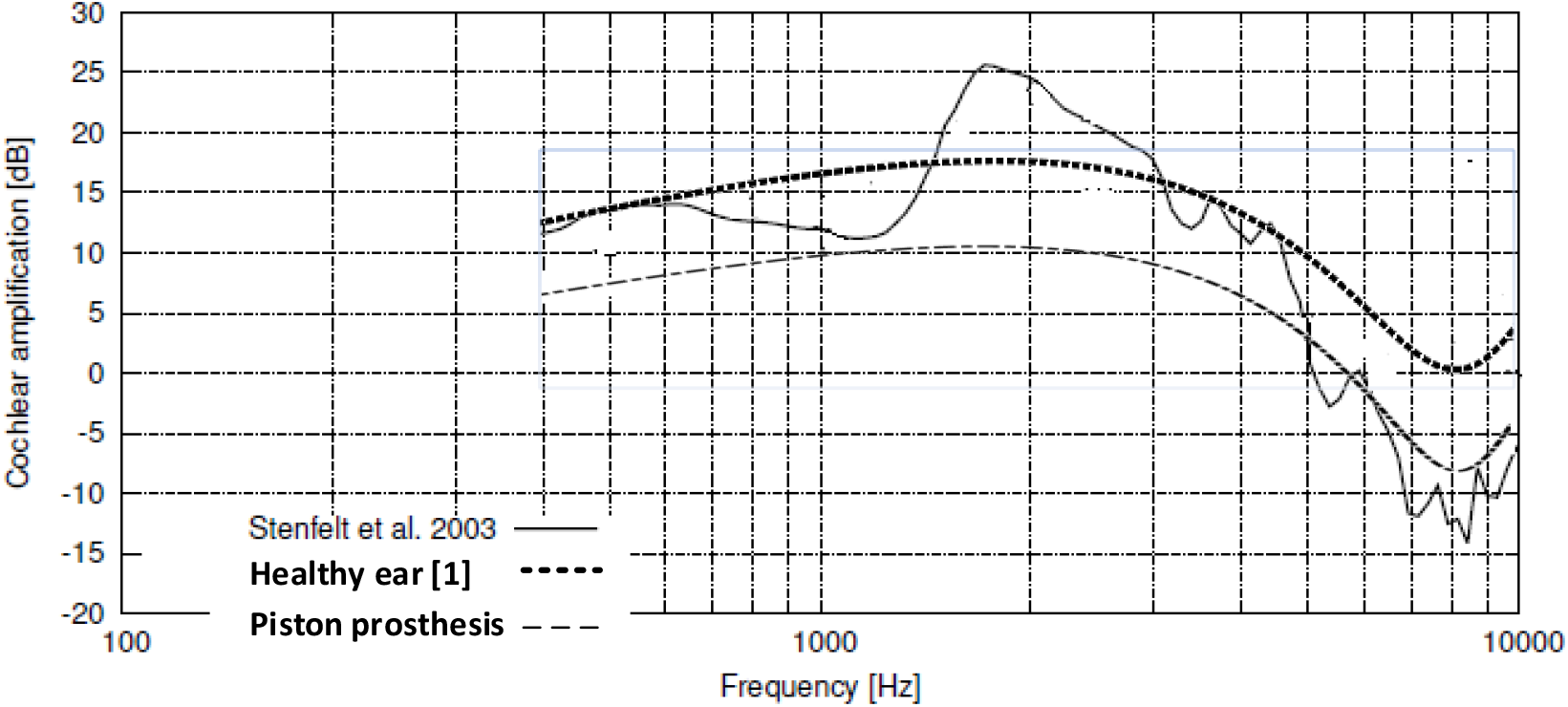
Level of cochlear amplification - test results and estimations for healthy ear and ear with piston prosthesis.

## 6. A modification of the piston prosthesis

### The principle of work of the piston in a guide tube

Note that to avoid the splitting of the sound wave in the cochlea, it is enough not to put the end of the piston into the cochlea. To do it, one can place the piston end, not inside the cochlea, but in a guide in the form of pipe ended with a funnel fixed in the hole made in the stapes footplate (Fig.8). To protect against too loud sounds, the tip of the piston should be at least 0.5 mm above the funnel (see [9]).

**Fig. 8.**
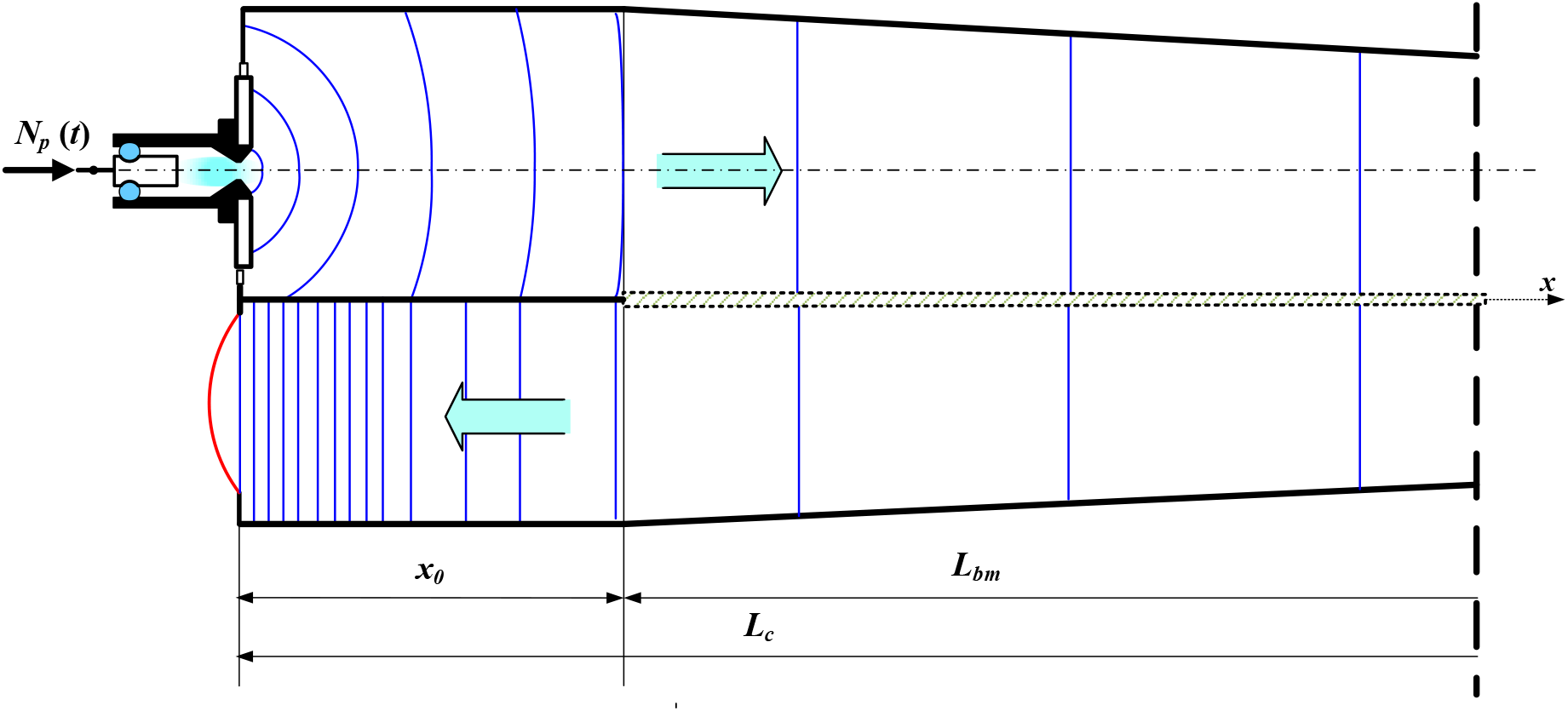
Wavefronts generated by vibrating piston prosthesis placed in the guide.

Now, the end of the piston makes the sound wave in the fluid filling this part of the guide that is directly connected to the cochlea. The double-cone shaped end of the guide focuses the sound wave in the hole leading to the cochlea, which can be taken as a point source of the sound. Spherical wave appears in the cochlea, which at some distance 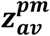 from the stapes footplate can be treated as a flat wave. To avoid the wave split effect, the outlet of the funnel should not stick out beyond the stapes footplate more than the radius of its edge rounding, i.e. 0.05 mm. Taking ***r***_***p***_ = **0** in Eq. (2), one can get

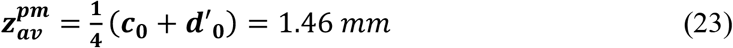

Recall the basilar membrane starts at the point ***x***_**0**_ = 2.6 *mm* from the stapes footplate. The point at which the sound wave becomes flat has the coordinate 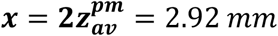. It is placed on the basilar membrane at the distance 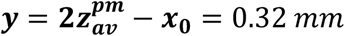 from its origin. This is 1% of the length ***L***_***bm***_ = 31.9 *mm* of the basilar membrane. One can find that at the point ***y*** the very high frequency of 1.9 kHz is received (see [Greenwood]). It means that the effect of the wavefront bending on the sound intensity in the cochlea can be neglected. Thus, one can state that the graph of the level of cochlear amplification (Fig. 7) for the ear with a modified prosthesis and for a healthy ear should be the same.

### A proposition of new piston prosthesis

The proposed prosthesis design is shown in Fig. 9a. In Fig. 9b, the partial axial section of the prosthesis shows its inside. The funnel outlet of the guide tube seen in Fig. 9b is filled with a physiological fluid just before the surgery.

**Fig. 9.**
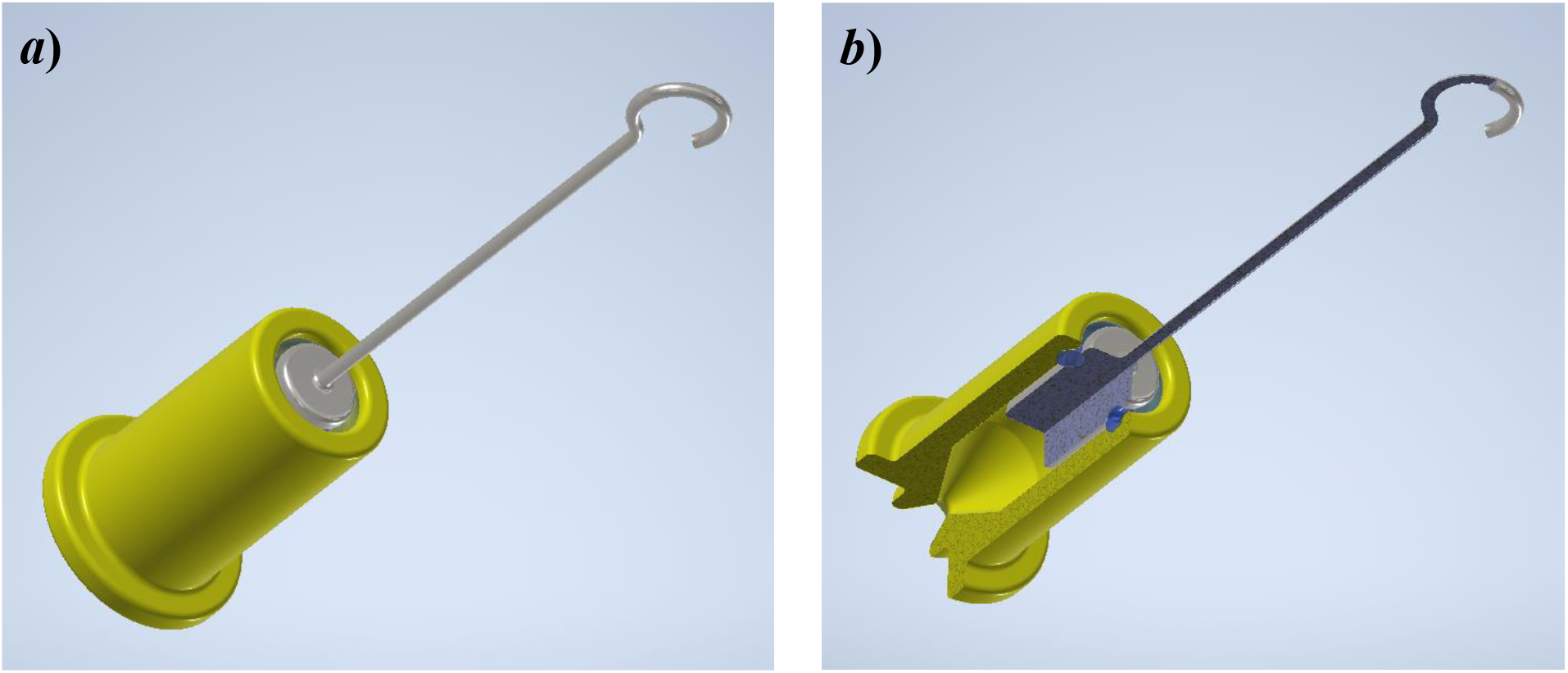
Modified piston prosthesis (a), its partial axial section (b).

The piston and the guide tube have a hollow (Fig. 10a,b) to allow the piston to seat in the tube on the O-ring formed of the elastic hydrogel (Fig 10c). Its stiffness is the same as the stiffness of the annular ligament in a healthy ear. This way enables to handle of the guide tube together with the piston.

**Fig. 10.**
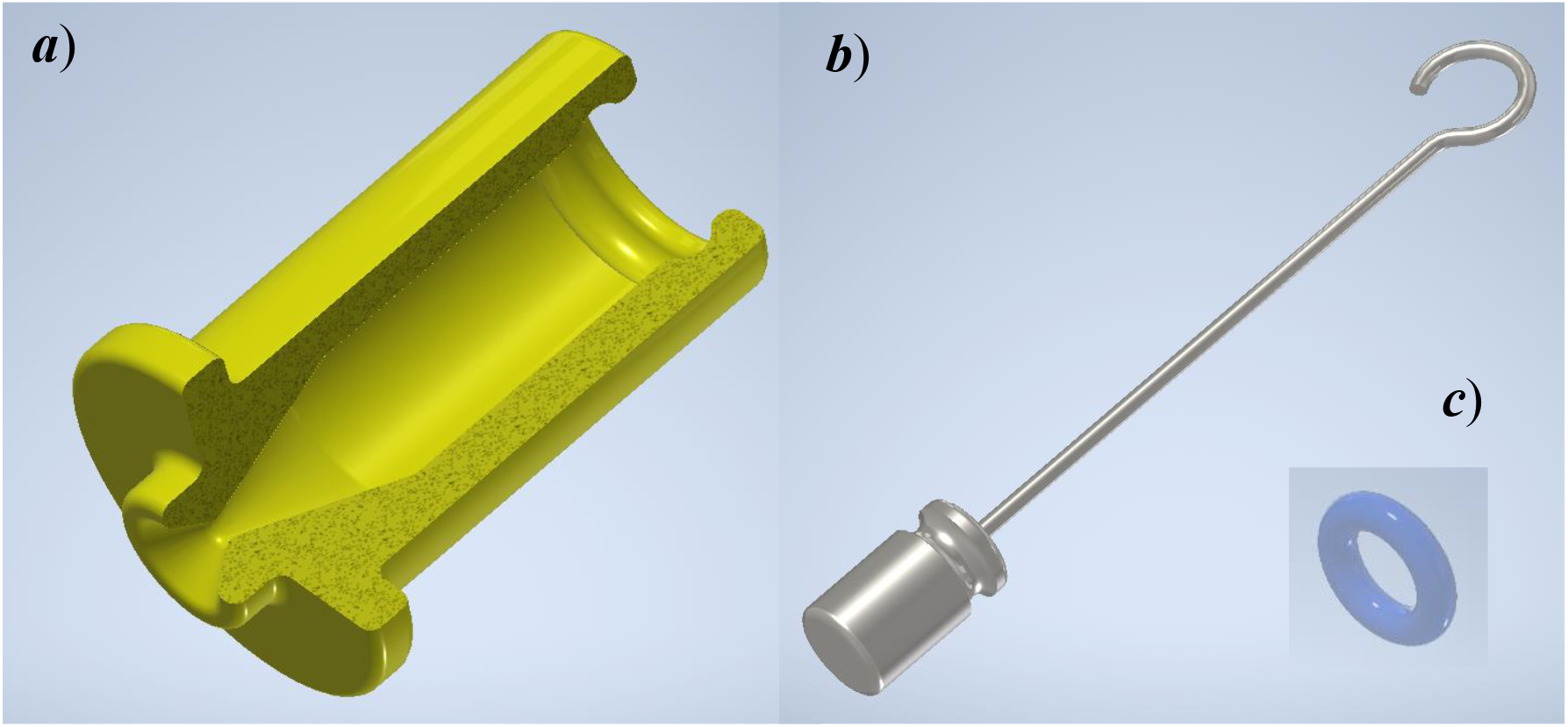
Elements of modified piston prosthesis: guide tube (a), piston (b), O-ring (c).

Let us assume that the O-ring with a radius ***r***_***O***_is sheared along the cylindrical walls of the piston and the guide with radii ***r***_***p***_ and ***r***_***g***_ (Fig.11).

**Fig. 11.**
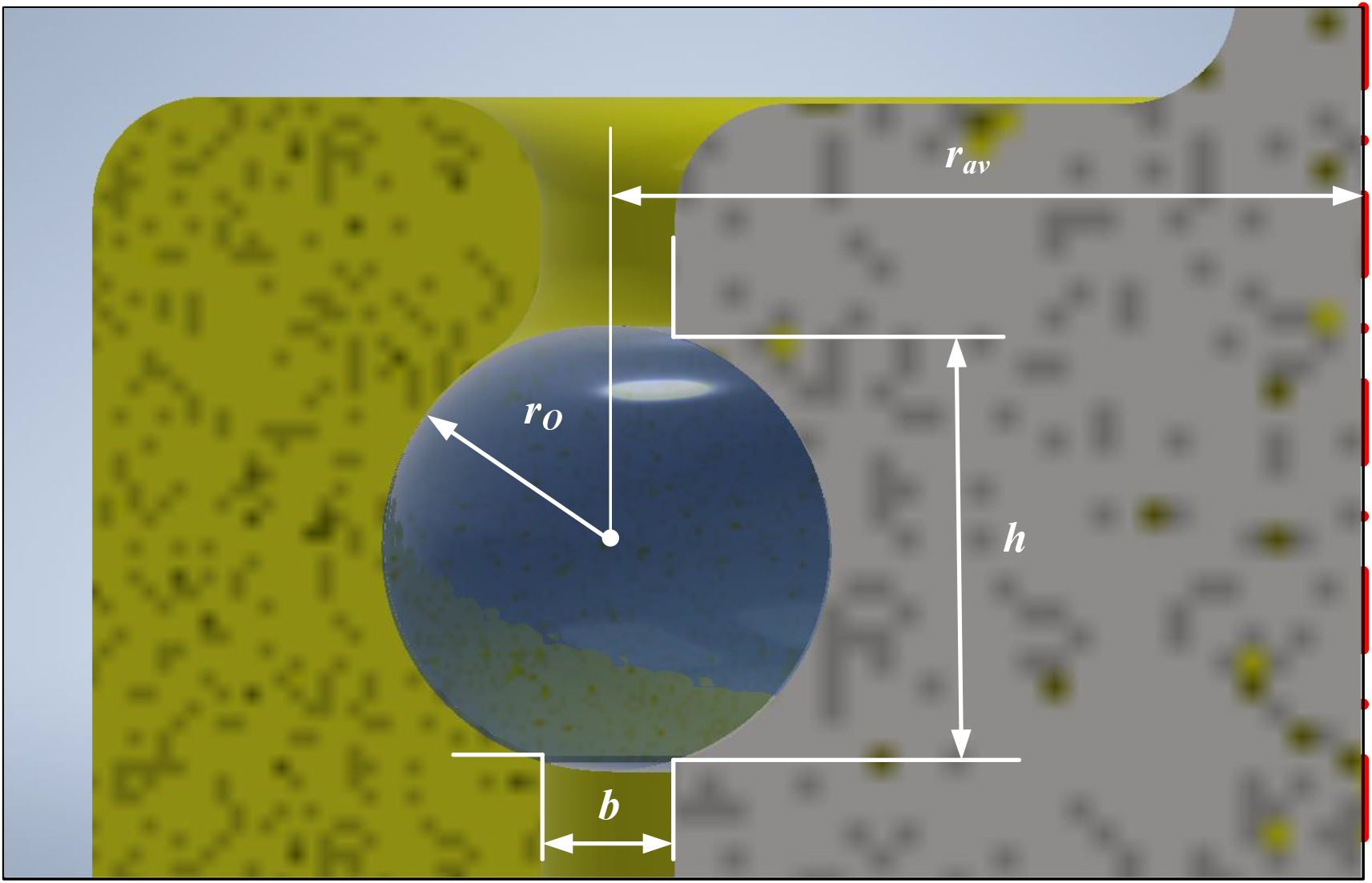
Details attachment of piston in the guide tube.

Denote by 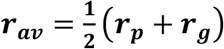 the average of these rays and by ***b*** = (***r***_***p***_ − ***r***_***g***_) and 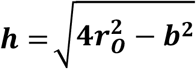, the width and the height of the shear cylindrical layer. One can find the Kirchhoff module ***G*** for the O-rig from the relation

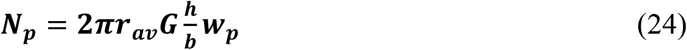

where ***N***_***p***_ = ***N***_***psf***_ = 1.4052 ⋅ 10^−6^ [*N*] is the force acting on the piston, and ***w***_***p***_ is the static amplitude of the piston. It is assumed that the last one is the same as that for the stapes footplate, and so ***w***_***p***_ = ***w***_***sf***_ = 1.17 ⋅ 10^−5^[*mm*]. Because 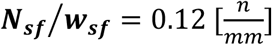, we have

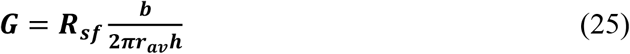

Taking ***r***_***aν***_ = 0.33 [*mm*], ***r***_***o***_ = 0.2 [*mm*], ***b*** = 0.06 [*mm*], one can get ***h*** = 0.19 [*mm*]. Then 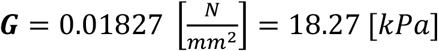 which is a typical value of the Kirchhoff module for silicone gels [10].

## 7. Discussion and conclusions

It was shown that too low stiffness of the piston suspension makes a drop of the amplitudes for the higher frequency vibrations. This disadvantage can be overcome by changing the piston suspension as it was shown in this work.

However, the main problem with the piston prosthesis is caused by the location of the piston end inside the cochlea. This disrupts a free propagation of the sound wave in the cochlea for the following reason. Around the edge of the piston end, which is inside the cochlea, the wavefront bends to form a toroidal surface. This divides the sound wave inside the cochlea. The main part of the toroidal surface is facing into the apex and forms a primary wave that runs along the cochlea axis. But a part of this surface is facing to the oval window and forms a backward wave. After the reflection from the stapes footplate, the backward waveforms a secondary wave that runs to the apex. The secondary wave follows the primary one with a small delay. This makes a drop in the intensity of the sound wave running in the cochlea and the formation of echo-like noises.

On the base of the simplified model of sound wave propagation in the cochlea, a graph of the level of cochlear amplification is made [5]. A comparison of this graph with the healthy ear graph shows a 5 dB drop. One can show that the curved front of the sound wave becomes flat over the basilar membrane. Thus, the curvature of the wavefront does not cause a drop in the sound intensity inside the cochlea.

To avoid the splitting of the sound wave in the cochlea, it is proposed to place the piston end, not inside the cochlea, but in a guide in the form of pipe fixed in the hole in the stapes footplate. In this case, the sound wave is formed under the piston end in the fluid filling the guide tube outside the cochlea. The funnel-shaped end of the guide focuses the sound wave in the cochlea inlet. The above idea is close to the concept of the chamber prosthesis. The last one gives a meaningful increase in the perception of high sounds [6].

The graph of the level of cochlear amplification for the piston in the guide tube is the same as in the case of the healthy ear. The shown details enable to make the new piston prosthesis simply and put it into practice.

## Acknowledgment

The author would like to thank Professor Monika Kwacz for a discussion and suggestions.

